# Allele-specific alternative polyadenylation links noncoding genetic variation to Alzheimer’s disease risk

**DOI:** 10.64898/2026.02.13.705798

**Authors:** Ryan M. Barney, Giovanni Quinones-Valdez, Alison J. King, Kofi Amoah, Weijian Wang, Xinshu Xiao

## Abstract

**Background:** Alternative polyadenylation (APA) is a crucial post-transcriptional mechanism generating isoform diversity in the nervous system. While genetic variants significantly influence gene expression, the extent to which they regulate 3’ UTR usage in the human brain remains underexplored. We aimed to characterize the landscape of allele-specific alternative polyadenylation (asAPA) and investigate its role in neurodevelopmental and neurodegenerative disorders.

**Results:** Analyzing 1,047 RNA-seq data from 293 Alzheimer’s disease (AD) and control donors across four brain regions, we identified 4,462 asAPA events involving 3,432 SNPs. We prioritized a core set of putative functional variants that drive consistent cis-regulatory effects across individuals. These functional SNPs are enriched for RNA-binding protein motifs, particularly those recognized by FMRP. In Fragile X Syndrome brains lacking FMRP, we observed widespread 3’ UTR shortening, with FMRP motifs enriched in the 3’ UTR extension regions of shortened transcripts, suggesting FMRP normally protects against proximal site usage. Integrating these data with population genetics, we found that asAPA SNPs significantly overlap GWAS risk loci for autism spectrum disorder (ASD), ADHD, and AD. Furthermore, comparing AD to control brains within our cohort revealed 77 asAPA genes exhibiting condition-specific shifts in allelic bias, affecting key synaptic genes including *CAMK2G*.

**Conclusions:** Our study uncovers a pervasive layer of cis-regulatory variation in the human brain that links noncoding genetics to transcript structure via RBP interactions. We identify FMRP as a key regulator of this process and demonstrate that asAPA provides a mechanistic bridge connecting genetic risk to neuronal pathology in both neurodevelopment and neurodegeneration.

## Introduction

Genome-wide association studies (GWAS) have linked thousands of common variants to complex traits and diseases, yet most of these variants fall in non-coding regions where the underlying mechanisms remain elusive [1]. One promising but underexplored avenue is post-transcriptional regulation, where variants can influence gene output well beyond changes in promoter or enhancer activities. In particular, alternative polyadenylation (APA) has emerged as a pervasive mechanism through which non-coding variants may act.

APA affects an estimated 70% of human genes, generating mRNA isoforms with distinct 3’ untranslated regions (3’ UTRs) [2,3]. Because 3’ UTRs harbor a repertoire of binding sites of microRNAs and RNA-binding proteins (RBPs), shifting polyadenylation sites can remodel mRNA stability, subcellular localization, and translation [4,5]. This regulation is particularly critical in neurons, where extended 3’ UTRs facilitate the transport of mRNAs to distal synaptic compartments for local translation [6,7]. Cis-regulatory elements such as polyadenylation signals and surrounding auxiliary motifs, together with trans-acting RBPs, orchestrate poly(A) site choice. Thus, genetic variants that disrupt these elements, or nearby RBP binding sites, have the potential to redirect APA and modulate gene function.

APA is dynamically regulated across tissues, developmental stages, and disease contexts, including neurodegeneration [8]. Prior transcriptome-wide studies have reported 3’ UTR shortening or lengthening in Alzheimer’s disease (AD), amyotrophic lateral sclerosis (ALS) and Parkinson’s disease [2]. While recent studies have begun to map the genetic architecture of these changes through alternative polyadenylation quantitative trait loci (aQTL) [9,10] a major challenge remains: standard aQTL mapping requires large sample sizes to achieve statistical power. This is a significant bottleneck for neurodegenerative research, where high-quality post-mortem brain cohorts are often limited in size. Thus, the specific contribution of genetic variants to APA dysregulation in AD remains incompletely understood.

Allele-specific APA (asAPA) analysis offers a powerful strategy to pinpoint genetically modulated APA events. By comparing the two alleles of heterozygous single-nucleotide variants (SNVs) within the same individual, asAPA controls for environmental and trans-acting effects, allowing for the sensitive detection of local genetic influences on poly(A) site choice [11]. This internal control makes asAPA uniquely suited for dissecting post-transcriptional regulation in post-mortem brain studies.

Here, we leverage asAPA to dissect genetic control of polyadenylation in the human brain and to explore its contribution to AD. Using RNA-seq data from four brain regions of the Mount Sinai Brain Bank (MSBB) cohort [12], we applied our allele-specific alternative mRNA processing (ASARP) pipeline to systematically catalog asAPA events, nominate putatively functional SNPs, and interrogate their impact on RBP binding, including the neuronal regulator FMRP. We further assess how allelic bias in APA differs between AD and control samples and evaluated the overlap of asAPA signals with aQTLs and GWAS loci for neurodegenerative and neurodevelopmental disorders. Together, our study offers a comprehensive view of allele-specific APA in the brain and highlights new connections between non-coding variation, transcriptomic regulation, and disease risk.

## Results

### Overview of asAPA landscape in the MSBB dataset

To characterize the landscape of genetically regulated polyadenylation in the human brain, we analyzed AD and controls brain RNA-seq data from the Mount Sinai Brain Bank (MSBB) [12] using the ASARP framework [11]. This approach distinguishes true asAPA from whole-gene expression differences by integrating allele-specific expression (ASE) with transcript structure. Specifically, we identified heterozygous SNVs located in alternatively processed 3’ UTRs that exhibit significant allelic imbalance, while ensuring that nearby SNVs in constitutive regions of the same gene show no imbalance. This criterion isolates cis-regulatory effects specific to the 3’ end. We note that the imbalanced SNVs associated with asAPA serve as markers (tag variants) for the allelic bias and are not necessarily the causal regulatory mutations.

After quality control and filtering (Methods), we analyzed 1,047 RNA-seq samples spanning four brain regions, frontal pole (BM10), inferior frontal gyrus (BM44), parahippocampal gyrus (BM36), and superior temporal gyrus (BM22), derived from 293 donors (Figure 1A). Applying ASARP across these samples, we identified a total of 4,462 unique asAPA events, involving 3,432 unique tag SNPs located in 1,040 distinct genes. To contextualize this, we tested a total of 26,221 distinct heterozygous SNPs in 3,228 genes with sufficient read coverage, indicating that approximately 13% of candidate SNPs and 32% of candidate genes showed significant allelic APA effects across all brain regions. While the cumulative number of significant events was large, individual samples typically contributed fewer than 20 events, possibly reflecting the strict coverage thresholds required to robustly detect allelic imbalance at APA sites (Supplementary Figure 1A; Methods).

**Figure 1.**
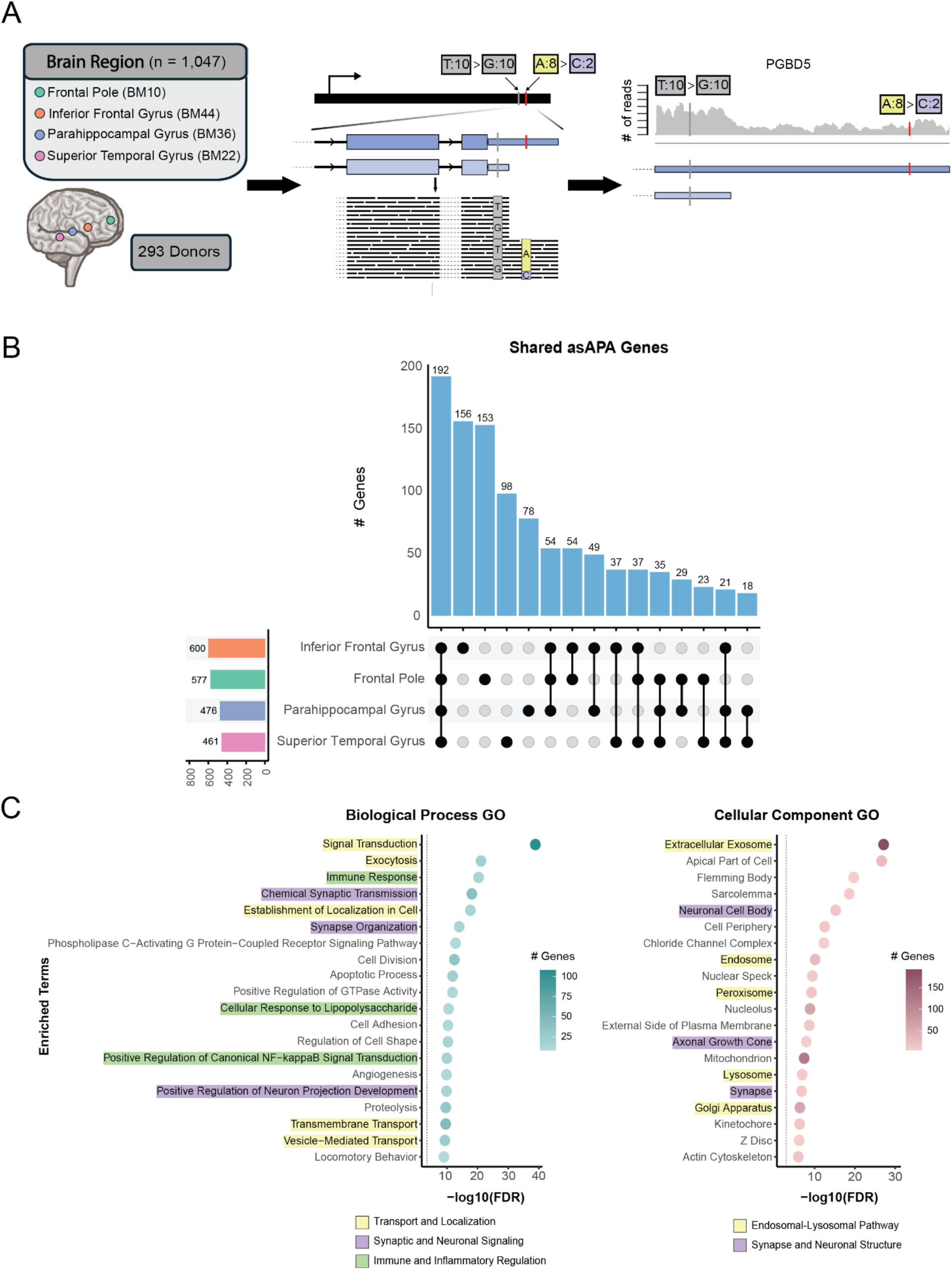
Overview of asAPA events in the human brain (A) Overview of the ASARP-based approach for detecting asAPA events. RNA-seq data from four brain regions across 293 donors (totaling 1,047 samples) were analyzed. Whole-genome sequencing (WGS) was used to identify heterozygous SNPs, and the ASARP method was applied to detect SNPs exhibiting allele-specific APA based on read imbalance in 3’ UTRs. As an example, in the PGBD5 gene, the A>C SNP was detected as the tag SNP exhibiting allelic bias (8:2 read counts) and located in the alternative 3’ UTR region; whereas the T>G SNP was the control SNP located in the core 3’ UTR and has no allelic bias (10:10 read counts). (B) Upset plot showing the overlap of asAPA genes across the 4 brain regions. The largest category consists of genes shared across all four regions. (C) Gene ontology (GO) enrichment analysis of asAPA-regulated genes. Left: Biological Process terms; Right: Cellular Component terms. Colors highlight broad categories as in the legends.

Assessing the overlap of asAPA *genes* across brain regions, we observed that the largest set consisted of genes shared by all four regions (Figure 1B). This trend still holds when restricting the comparison to genes testable in all brain regions (Supplementary Figure 1B). In contrast, most asAPA-associated *SNPs* were unique to individual brain regions (Supplementary Figure 1C, 1D). This divergence, broad overlap at the gene level versus specificity at the SNP level, suggests that many identified variants act as tag SNPs rather than causal drivers. While the specific tag SNP capturing a haplotype may vary across individuals (and thus brain regions), they likely index the same underlying cis-regulatory event. This interpretation is supported by the high similarity of asAPA gene sets between pairs of brain samples (Supplementary Figure 1E), consistent with different SNPs in linkage disequilibrium (LD) capturing the same regulatory effect.

To investigate the functional relevance of these asAPA genes, we performed Gene Ontology (GO) enrichment analysis using the full set of 1,040 genes and a background of tested but non-asAPA genes (Methods). This analysis revealed strong enrichment for biological processes involved in intracellular trafficking, vesicle transport, and subcellular localization (Figure 1C, left, yellow-highlighted). Genes involved in neuronal signaling and synaptic organization (Figure 1C, left, purple-highlighted), and immune and inflammatory processes (Figure 1C, left, green-highlighted) are also enriched. Concordantly, cellular component terms were enriched for vesicle-associated compartments such as endosome, lysosome, Golgi apparatus, and peroxisome (Figure 1C, right, yellow), reinforcing the involvement of APA-regulated transcripts in membrane trafficking pathways. While this analysis is not disease-specific, the convergence on endosomal-lysosomal and trafficking-related components is particularly intriguing given their central role in maintaining neuronal homeostasis and their disruption in various neurodegenerative diseases, including AD [13].

### Identifying putative functional asAPA SNPs

While the majority of asAPA-associated SNPs exhibit region-specificity, this likely reflects LD rather than biological variation. To prioritize SNPs with potential regulatory impact, we applied a concordance-based analysis [14] that evaluates whether a SNP consistently predicts 3’ UTR isoform usage across all heterozygous individuals (Methods). This approach leverages the expectation that truly functional cis-regulatory variants drive concordant APA shifts in heterozygous individuals across the cohort, thereby filtering out linkage-driven associations.

Applying this approach in each of the four brain regions, we identified a total of 167 unique high-concordance asAPA SNPs mapping to 67 distinct genes. In stark contrast to the initial broad set of variants (Supplementary Figure 1C), this functionally enriched subset showed a high degree of sharing across brain regions (Figure 2A). Indeed, the largest category of putatively functional SNPs (74 SNPs) and genes (24 Genes) were those shared by all four brain regions. For example, the *PRXL2A* locus (Supplementary Figure 2A) illustrates a high-concordance SNP that is active across 4 brain regions. This convergence suggests that by filtering for concordance, we have captured a core set of cis-regulatory mechanisms that are conserved across the brain. Consequently, we defined this set as “putatively functional SNPs” and utilized them for downstream characterization of post-transcriptional regulation.

**Figure 2.**
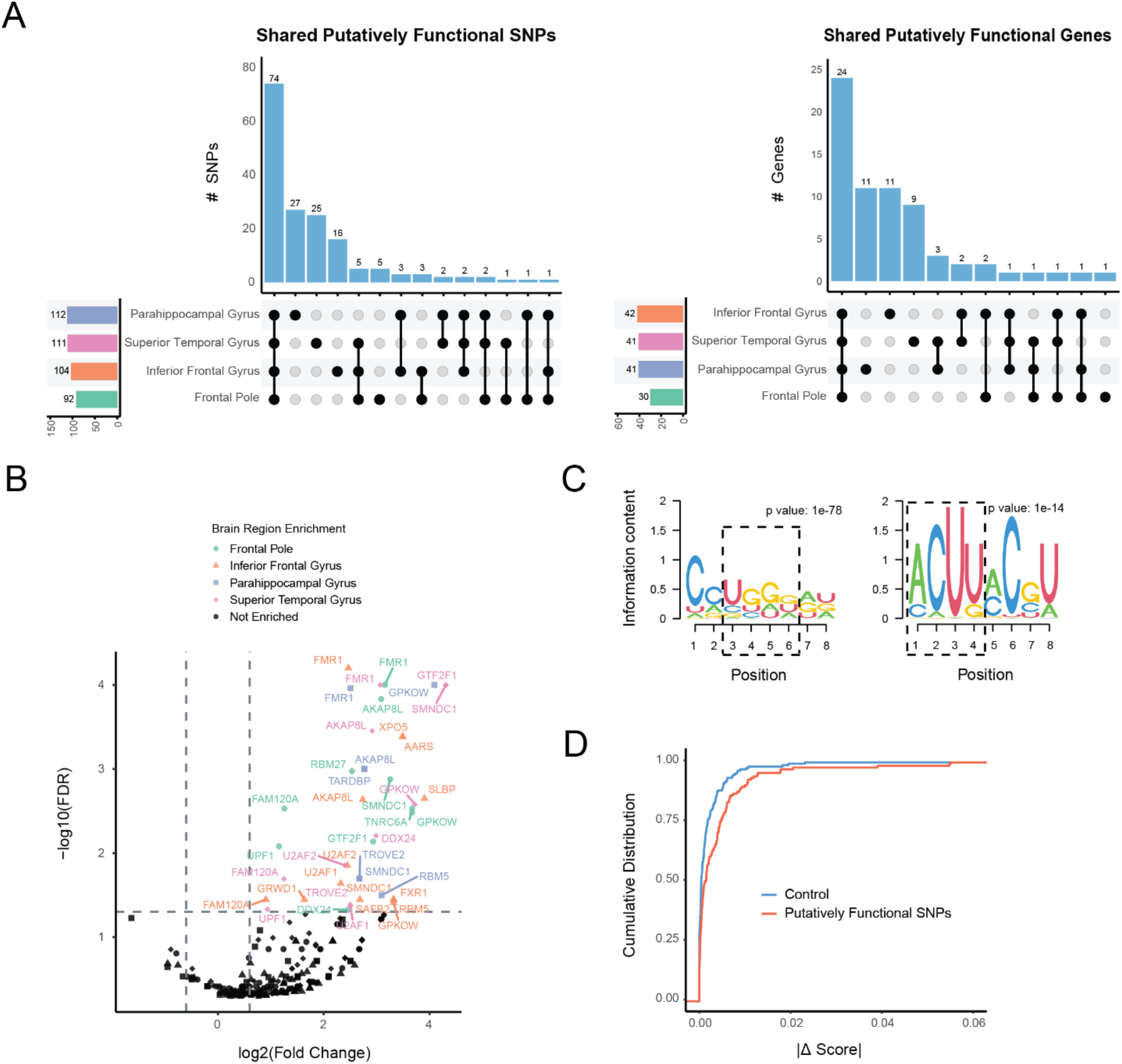
Identification of putatively functional asAPA SNPs and their association with RBPs. (A) Upset plots showing overlap of putatively functional asAPA SNPs (left) and their host genes (right) across the four brain regions. (B) Enrichment of RBPs whose eCLIP binding peaks overlap regions harboring putatively functional asAPA SNPs. Fold change was calculated as the ratio between the proportion of genes harboring putatively functional asAPA SNPs bound by each RBP and the proportion of genes harboring control SNPs bound by each RBP, where control SNPs were those tested by ASARP but not significant. P values were obtained from bootstrapping the control genes to generate a null distribution and corrected with the Benjamini-Hochberg method. Genes with FDR < 0.05 and fold change >1.5 are highlighted. (C) Two motifs from *de novo* motif identification by HOMER2 around putatively functional asAPA SNPs. Motifs resemble previously reported FMRP binding preferences, including UGGA and ACUU. Hypergeometric test p-value comparing motif frequency in higher expressed alleles to lower expressed alleles is shown. (D) Cumulative distribution of absolute binding score differences between alleles for *FMR1*, as predicted by DeepRiPe. Putatively functional asAPA SNPs (red) show significantly greater predicted allele-specific impact on FMRP binding compared to control SNPs (blue) (Kolmogorov-Smirnov test, FDR = 0.006).

### Putatively functional asAPA SNPs alter RNA-protein interaction landscape

Having prioritized a set of putatively functional asAPA SNPs, we next investigated their potential regulatory consequences, specifically, how they may alter interactions with RBPs, which are known to regulate polyadenylation. Using eCLIP data from ENCODE [15], we performed an enrichment analysis to identify RBPs that bind to putatively functional asAPA SNPs more frequently than control SNPs (Methods). This analysis revealed a significant enrichment of binding events for several RBPs across brain regions (Figure 2B). Notably, FMRP (encoded by the *FMR1* gene) emerged as a consistently significant hit in every region, supporting a potential role for *FMR1* in modulating APA across the brain.

To complement the eCLIP-based analysis, we used HOMER2 for de novo motif discovery to detect sequence-specific drivers (Methods). We compared motifs present on alleles with higher RNA-seq read counts (“preferentially expressed”) versus those with lower counts and matched the motifs to RBPs via the RNA Bind-n-Seq (RBNS) data [16]. This analysis successfully recovered FMRP-associated recognition motifs on the preferentially expressed alleles, including the canonical UGGA and ACUK binding sequences [17]. We also observed significant motif enrichment for other key neuronal regulators, including TIA1, CPEB1, and ELAVL4 (Supplementary Figure 2B), reinforcing the link between these functional SNPs and neuronal RNA regulation [18,19,20].

Finally, we used DeepRiPe, a deep learning-based framework, to predict the impact of sequence variants on RBP binding (Methods). Comparing the functional asAPA SNPs to matched controls, we found that the functional variants were predicted to cause significantly greater disruption to RBP binding profiles for several RBPs (Supplementary Figure 2C). Consistent with our eCLIP and motif findings, the predicted binding scores for FMRP were significantly altered by these functional SNPs (Figure 2D), providing computational validation that these genetic variants likely modulate FMRP-mediated regulation.

### FMRP is associated with 3’ UTR regulation in the human brain

Given FMRP’s consistent enrichment in our RBP analyses, including eCLIP binding, motif enrichment, and allelic binding predictions, we next investigated whether it plays a regulatory role in 3’ UTR processing. To address this, we analyzed RNA-seq data from postmortem frontal cortex tissues of individuals with Fragile X Syndrome (FXS), a condition caused by epigenetic silencing of the *FMR1* gene and consequent loss of FMRP expression [21]. These data included two cohorts: one consisting of two FXS patients and two controls, and a second consisting of two FXS patients and two premutation carriers [22]. Together, these samples provide a human disease model to examine FMRP-dependent APA dynamics in the brain.

Using annotated polyadenylation sites from polyA_DB [23], we quantified 3’ UTR isoform usage to identify systematic shifts in FXS (Methods). In both cohorts, we observed widespread shifts in APA usage, with hundreds of transcripts exhibiting either shortening or lengthening in FXS relative to comparison groups (Figure 3A). These changes could reflect a combination of direct and indirect consequences of FMRP loss.

**Figure 3.**
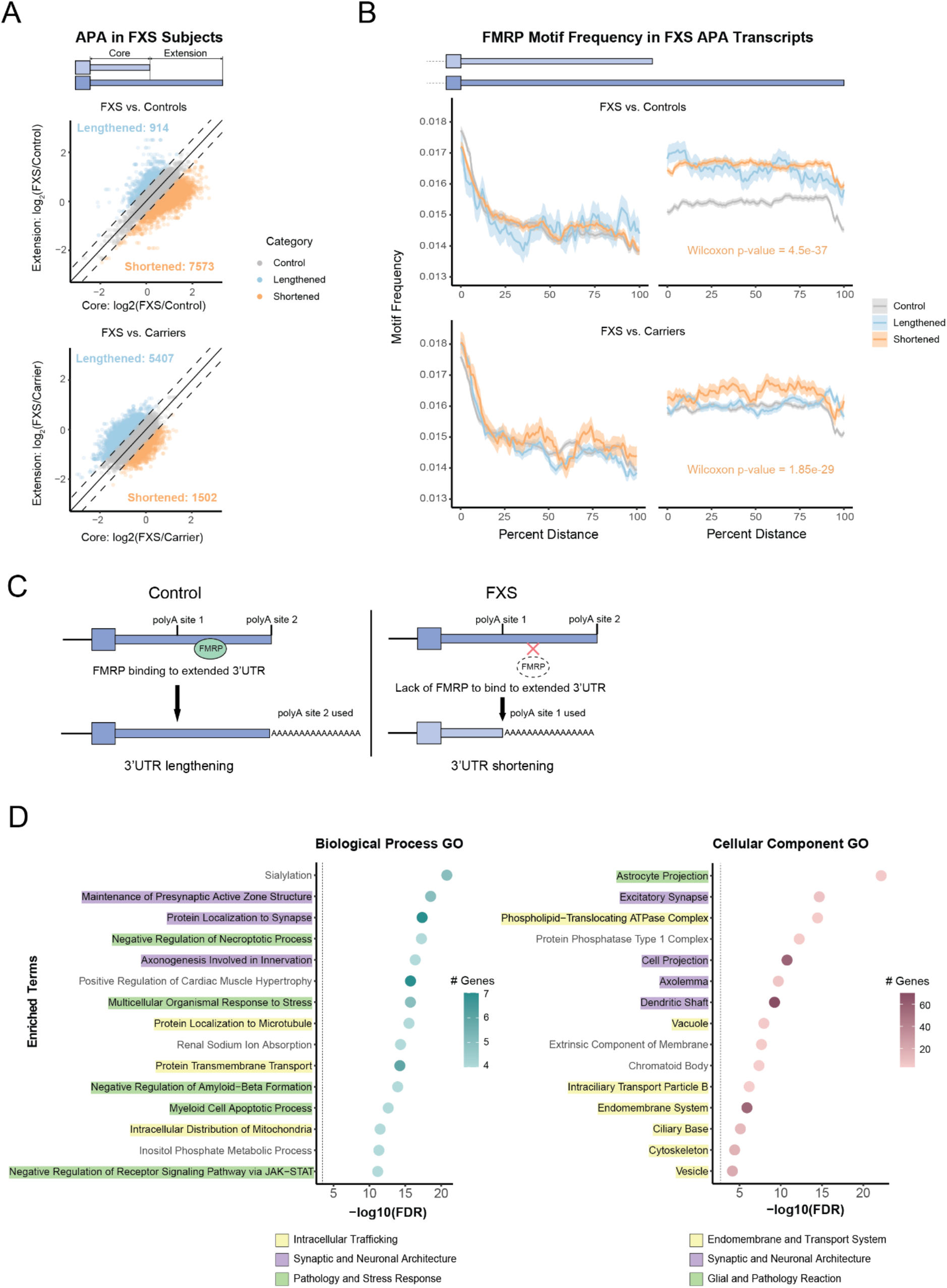
Loss of FMRP in FXS patients is associated with APA changes (A) Scatter plots comparing APA isoform usage in FXS versus control (top) and FXS versus carrier (bottom) brain samples. Each point represents a transcript; axes show log_2_ fold change between FXS and controls (or carriers) in the usage of core or extension regions, where usage is calculated as the average coverage of a 400bp window upstream of the end of the core or extension region. Transcripts are colored by the direction of APA change (using a threshold of 0.5 for |log2 (FXS/control)| or |log2 (FXS/carrier)|: lengthened (blue) or shortened (orange) in FXS relative to comparison group. (B) FMRP motif frequency (mean ± standard error of the binomial proportion) in the core and extension regions of 3’ UTRs for transcripts showing 3’ UTR shortening (orange), lengthening (blue), or no change (gray) in FXS. Top: FXS vs. control; Bottom: FXS vs. carrier. Consistent to both cohorts, significantly higher motif density was observed in the extended regions of transcripts shortened in FXS. (C) Model illustrating FMRP’s proposed role in APA regulation. Under normal conditions (left), FMRP binds in the extension regions to promote 3’ UTR lengthening. In FXS (right), FMRP loss leads to increased usage of proximal sites and 3’ UTR shortening. (D) GO enrichment analysis of genes with shortened 3’ UTRs in FXS. Left: Biological Process terms; Right: Cellular Component terms; Colors highlight broad categories as in the legends.

To specifically assess the role of FMRP in regulating these APA changes, we examined the occurrence of known FMRP motifs within lengthened and shortened 3’ UTRs (Methods). We found that transcripts exhibiting 3’ UTR shortening in FXS were more enriched for FMRP motifs in the extension regions of the 3’ UTRs compared to control UTRs (Figure 3B). This pattern was consistent across both cohorts, supporting a model in which FMRP normally binds in the extended 3’ UTR region to promote distal polyA site usage or inhibit proximal site recognition. In the absence of FMRP, this regulatory influence is lost, leading to preferential use of proximal sites and resulting in 3’ UTR shortening (Figure 3C).

To explore the functional relevance of transcripts exhibiting APA changes in FXS, we performed GO enrichment analysis on genes with significantly shortened 3’ UTRs. Biological Process analysis (Figure 3D, left) revealed enrichment for terms associated with synaptic maintenance (e.g., Maintenance of Presynaptic Active Zone Structure, Protein Localization to Synapse) and stress response signaling (e.g., Multicellular Organismal Response to Stress, Negative Regulation of Amyloid-Beta Formation). Complementing these functional findings, Cellular Component analysis (Figure 3D, right) localized these changes to specific neural architectures, including excitatory synapses, dendritic shafts, and astrocyte projections. Notably, the strong enrichment for endomembrane and vesicular systems mirrors the pathways identified in our global asAPA analysis (Figure 1), supporting that FMRP is a key driver of 3’ UTR regulation in neuronal trafficking and synaptic communication networks.

### asAPA SNPs explain aQTL and GWAS signals

To assess whether asAPA is associated with broader patterns of genetic variation, we next asked whether asAPA SNPs overlap known genetic signals from aQTL and GWAS. While these large-scale studies have uncovered many noncoding variants linked to neurodevelopmental and neurodegenerative traits, the mechanisms through which these variants act remain incompletely understood. We hypothesized that genetically regulated APA events captured by our asAPA framework could mechanistically explain a portion of these associations.

We first compared our set of asAPA-associated genes to genes linked to aQTL in GTEx brain tissues [9]. Approximately half of the asAPA genes (540 of 1,040) overlapped with reported aQTL genes (Figure 4A), indicating a substantial intersection between allele-specific APA and population-level APA variation. At the SNP level, asAPA-associated variants were significantly more likely than control SNPs (defined as tested but nonsignificant asAPA SNPs) to overlap reported aQTLs (Supplementary Figure 3A). In addition, asAPA SNPs displayed stronger statistical significance in the aQTL dataset compared to matched controls (Supplementary Figure 3B). This enrichment was also pronounced at the gene level (Figure 4B). In a quantile-quantile (QQ) analysis comparing aQTL p-values of asAPA genes against controls, asAPA genes showed a notable deviation from expectation, consistent with stronger enrichment for aQTL signals.

**Figure 4.**
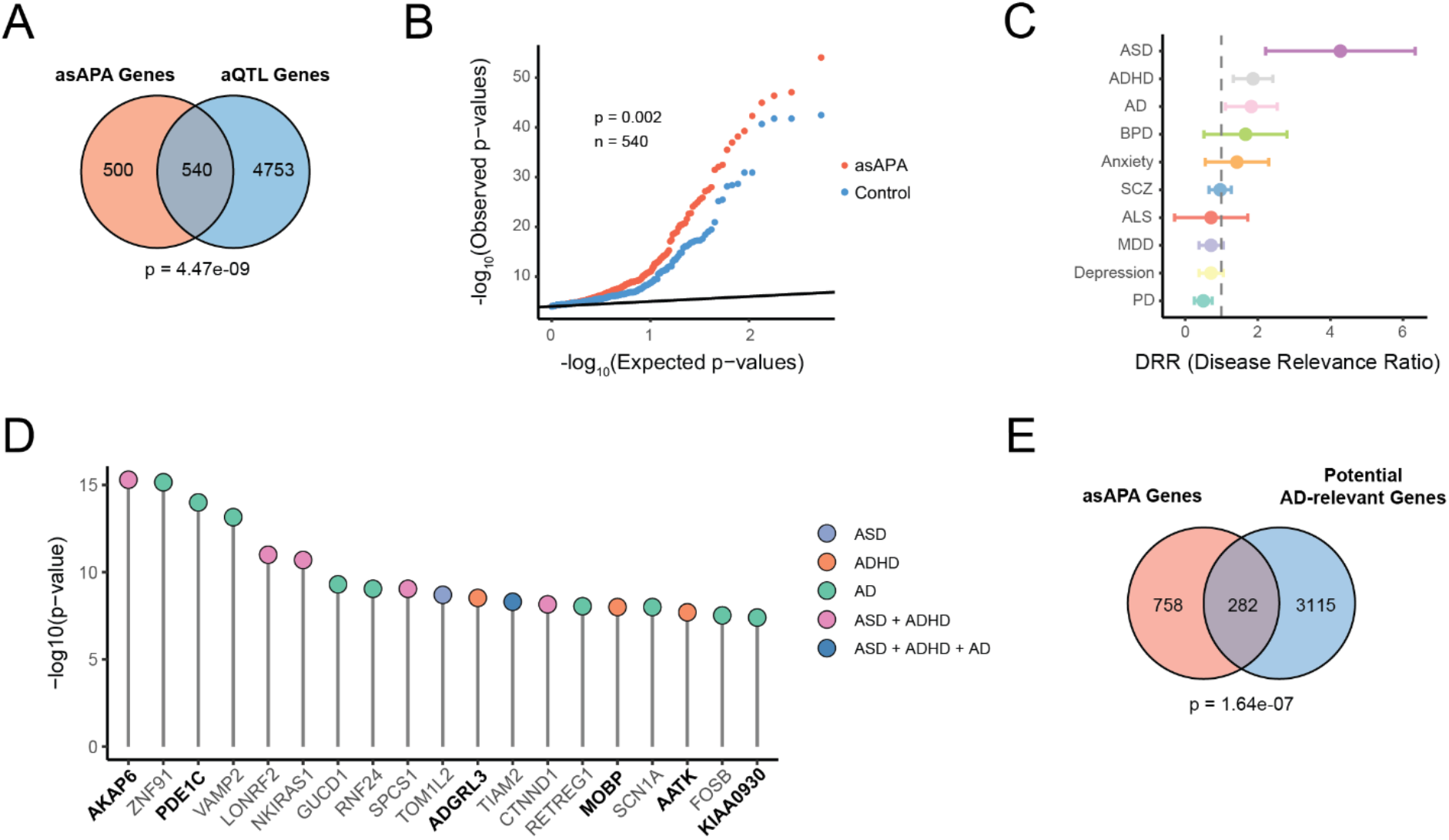
asAPA SNPs overlap with aQTL and GWAS signals relevant to brain disorders (A) Venn diagram showing the total number of asAPA genes overlapping reported aQTL genes from GTEx brain tissues. Statistical significance was assessed using a Chi-square test comparing overlap with a control gene set defined as genes containing exclusively SNPs that were testable by ASARP but not significant. (B) QQ plot of aQTL p-values for putatively functional asAPA genes (red) compared to the same control genes defined in (A) (blue), where the minimum p-value is taken if multiple exist for a given gene. Kolmogorov-Smirnov test p value is shown. (C) Disease relevance ratio (DRR) comparing the frequency of GWAS overlap among asAPA SNPs to that among matched control SNPs (testable by ASARP but not significant) for ten neurodevelopmental and neurodegenerative disorders. A DRR > 1 (dashed line) indicates enrichment. (D) GWAS p-values for asAPA SNPs overlapping GWAS loci, colored by the associated disease(s). Genes containing SNPs that directly overlap a GWAS index SNP (not just in LD) are bolded. (E) Venn diagram showing overlap between asAPA genes and previously reported AD-relevant genes. Chi-square test as in (A).

To evaluate whether genetically regulated APA is associated with common disease risk, we assessed the overlap of asAPA SNPs with GWAS loci from ten neurodevelopmental and neurodegenerative disorders [1,24]. A SNP was considered overlapping if it was in LD (r^2^ ≥ 0.6) with a GWAS index SNP and located within 200 kb. To quantify enrichment, we calculated a disease relevance ratio (DRR), comparing the frequency of GWAS overlap among asAPA SNPs to that among matched control SNPs. A DRR > 1 indicates enrichment. Notably, we observed elevated DRRs for ASD, attention-deficit/hyperactivity disorder (ADHD), and AD, suggesting that APA regulation may be functionally linked to the genetic risk architecture of these conditions (Figure 4C). Representative genes harboring GWAS-overlapping asAPA SNPs associated with ASD, ADHD, or AD, including *AKAP6, PDE1C* and *ADGRL3*, are shown in Figure 4D. The ASD signal is particularly intriguing in light of our earlier findings implicating FMRP (the loss of which causes FXS, a leading genetic cause of autism) as a master regulator of APA.

To complement these variant-level associations, we tested whether asAPA-regulated genes were enriched for previously reported AD-relevant genes derived from transcriptomic studies of human brain tissue [25,26]. We observed a significant overlap, with 282 asAPA genes also appearing in the AD-associated gene set (Figure 4E), supporting the hypothesis that APA regulation may contribute to molecular processes altered in AD.

### asAPA profiles are altered in Alzheimer’s disease

Having established that asAPA events are enriched at genetic loci associated with AD, we next investigated whether allele-specific APA regulation differs between AD and control brains. To classify samples, we used a composite neuropathology-based grouping that integrated CERAD scores, Braak stages, and amyloid-beta burden, following previously described criteria (Methods) [27]. Differential allelic expression at asAPA SNPs was then assessed using REDIT-LLR, a beta-binomial framework originally developed for detecting condition-specific RNA editing differences (Figure 5A) [28]. We confirmed the applicability of this approach for allele-specific APA analysis (Methods, Supplementary Figure 4A).

**Figure 5.**
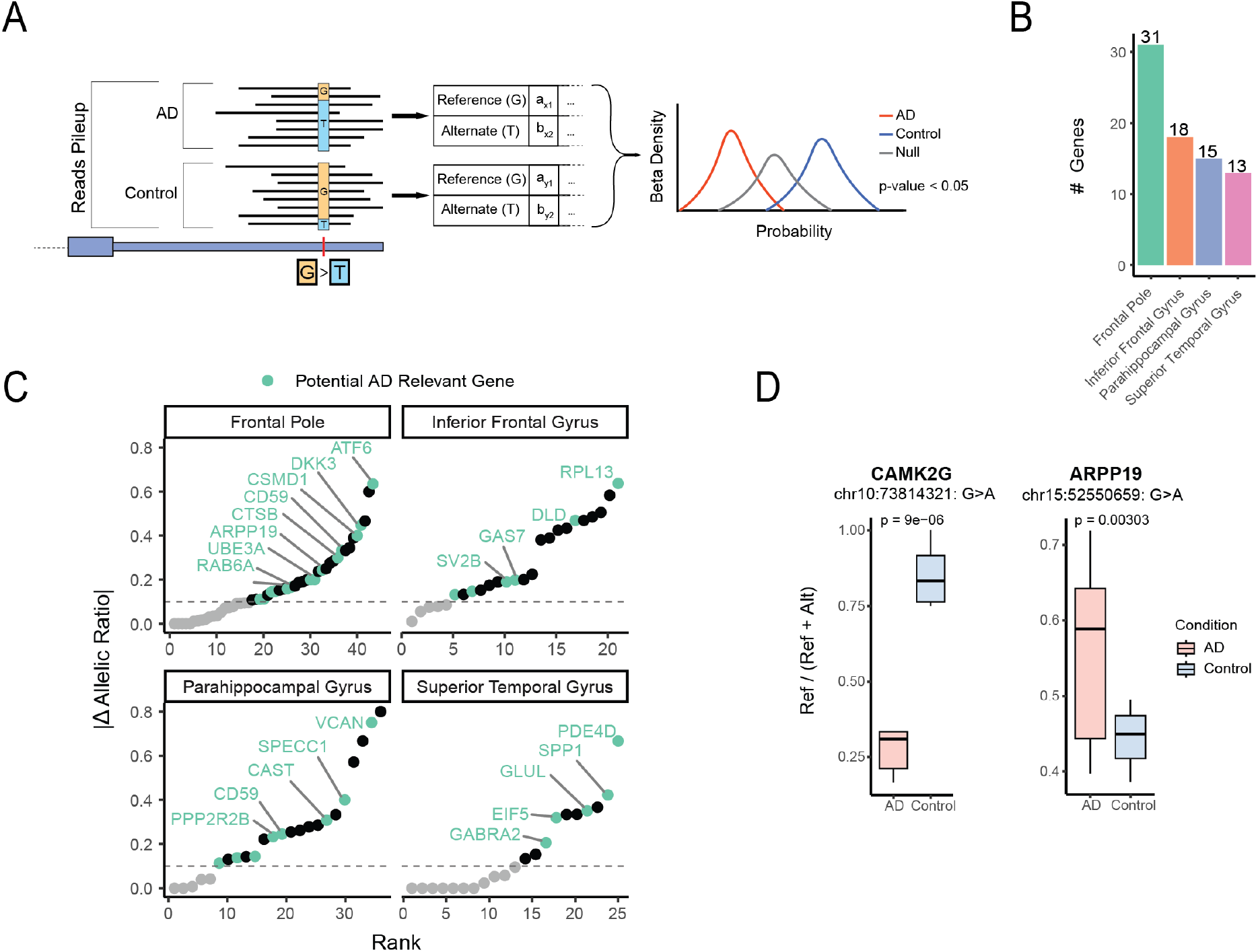
Condition-specific allelic APA changes in AD (A) Schematic of the REDIT-LLR framework applied to asAPA SNPs. Allelic counts for reference and alternative alleles in AD and control groups were modeled using a beta-binomial distribution to test for differential allelic bias between conditions (Methods). (B) Number of genes with significant differential allelic APA events in each brain region. (C) Ranked plots showing the magnitude of allelic shift (|Δ allelic ratio|) between AD and controls across brain regions. Genes previously implicated in AD are labeled. (D) Examples of differential asAPA events. *ARPP19* shows increased reference allele usage in AD, while *CAMK2G* shows increased alternative allele usage in AD. Both genes are involved in neuronal signaling and function. Differential allelic usage p-value was calculated using REDIT-LLR (Methods).

Applying REDIT-LLR across brain regions to asAPA SNPs that passed quality filters (Methods), we identified 77 significant genes (including 82 unique SNPs) that exhibit differential allelic bias between AD and control groups (Figure 5B). These findings suggest that disease status is associated with condition-specific shifts in APA regulation at the cis level. The direction of this shift was locus-dependent, with no global systematic bias toward either the reference or alternative allele (Supplementary Figure 4B). Notably, several of the genes showing differential allelic bias have been previously implicated in AD biology [25,26] (Figure 5C).

To illustrate these disease-specific differences, we highlight two representative examples (Figure 5D). In *CAMK2G*, a calcium/calmodulin-dependent kinase critical for synaptic plasticity [29,30], the allelic ratio shifted significantly in AD samples, reflecting increased usage of the alternative allele. In contrast, *ARPP19*, a cAMP-regulated phosphoprotein implicated in neurodevelopment and mitotic regulation [31,32], showed a shift toward greater usage of the reference allele in AD. Both genes are central to neuronal signaling and plasticity, raising the possibility that APA changes at these loci may contribute to altered neuronal function in AD.

## Discussion

Our study presents a systematic analysis of allele-specific alternative polyadenylation (asAPA) in the human brain, revealing a widespread but underexplored layer of post-transcriptional regulation. By leveraging genotype-resolved RNA-seq data across four brain regions, we identified over 4,400 asAPA events that converge on pathways involved in vesicle trafficking, endosomal-lysosomal function, and neuronal signaling, processes essential for synaptic maintenance and neuronal homeostasis [33]. These results suggest that genetic variation influencing 3’ UTR usage is not an isolated phenomenon but a pervasive mechanism shaping transcript diversity and regulation in the brain.

A particularly compelling insight from our analysis is the repeated convergence on FMRP. Putatively functional asAPA SNPs, those with consistent allelic effects across individuals, were enriched for predicted disruptions of RNA-binding protein interactions, with FMRP emerging as a top candidate. FMRP is a translational regulator implicated in FXS, a monogenic form of autism spectrum disorder [34,35,36,37], and Fragile X-associated tremor/ataxia syndrome (FXTAS), a late-onset neurodegenerative condition with cognitive and motor symptoms overlapping those seen in AD [38,39,40]. This overlap raises the possibility that FMRP-regulated APA could be a shared molecular feature relevant to both neurodevelopment and neurodegeneration.

Consistent with this idea, we observed global 3’ UTR remodeling in FXS brains, with significant enrichment of FMRP motifs in the extended UTR regions of transcripts shortened in FXS. These observations support a model in which FMRP binding normally “protects” the extension region to suppress proximal site usage, and its loss shifts the transcriptome toward shorter isoforms. Many affected genes were enriched for synaptic function and membrane transport, processes that are central to synaptic vesicle recycling, receptor turnover, and proteostasis [41]. Given that endosomal-lysosomal dysfunction is a hallmark of AD pathology [42], this model suggests a potential mechanistic bridge linking post-transcriptional misregulation in FXS, FXTAS, and AD.

Integrating asAPA with population-level aQTL data [9] revealed substantial overlap at both the gene and SNP level, suggesting that allele-specific APA may underlie a portion of population-level APA variation. We also observed significant enrichment of asAPA SNPs in GWAS loci [1,24] for ASD, ADHD, and AD, nominating APA as a possible molecular interface between noncoding genetic variation and disease-relevant pathways. Notably, many genes harboring GWAS-overlapping asAPA SNPs intersect with known AD-relevant genes [25,26]. In addition, the strong enrichment in ASD is particularly intriguing given that mutations in *FMR1* represent a monogenic cause of ASD via FXS, suggesting the GWAS signal we observe may, in part, be explained by the relationship between asAPA SNPs and FMRP-mediated regulation. While these associations do not establish causality, they provide a framework for investigating how noncoding variants might modulate isoform usage and downstream regulatory potential.

Our direct comparison of AD and control brains revealed 82 asAPA SNPs exhibiting significant condition-specific shifts in allelic bias. These changes may represent disease-associated perturbations in APA regulation, possibly through altered RBP activity or changes in the cis-regulatory landscape. Two illustrative examples, *CAMK2G* and *ARPP19*, are key regulators of synaptic plasticity and phosphoregulation, respectively [29,30,31,32]. Although further validation is needed, these findings support a model in which APA rewiring contributes to the synaptic dysfunction that is a central pathological feature of AD.

Our study also presents several methodological strengths and limitations. By applying a concordance-based framework, we were able to prioritize a subset of putatively functional asAPA SNPs, those with consistent genotype-APA relationships across individuals, thereby increasing the specificity for identifying cis-regulatory variants with likely biological impact. This within-dataset consistency strengthens causal inference and complements population-level association approaches. However, the power of this analysis remains constrained by sample size and the need for sufficient read coverage at heterozygous sites, which limits detection sensitivity. Additionally, the use of bulk RNA-seq data masks cell-type-specific APA regulation. Future studies incorporating single-cell sequencing technologies will be essential to map asAPA regulation with greater cellular specificity, particularly in the context of disease.

In summary, by bringing allele-specific APA into the framework of post-transcriptional regulation in the brain, our work highlights how noncoding variation can shape not only expression levels but also transcript structure and regulatory potential. This study provides a foundation for dissecting how APA intersects with RBPs like FMRP to influence neuronal function, and points to asAPA as a promising axis for understanding, and potentially intervening in, the molecular pathways underlying neurodevelopmental and neurodegenerative disorders.

## Methods

### Data preprocessing and asAPA identification

We obtained aligned RNA-seq BAM files and WGS-derived variants of the MSBB samples from synapse.org (syn3159438), derived from postmortem human brains across AD neuropathological stages [12]. RNA-seq data from four regions were available: frontal pole (BM10), inferior frontal gyrus (BM44), parahippocampal gyrus (BM36), and superior temporal gyrus (BM22).

The following quality control steps were performed: (1) BAM files flagged by MSBB for metadata mismatches were removed. (2) For donors with multiple sequencing replicates, only the most recent replicate was retained. (3) Samples lacking genotype data were excluded, as they were not suitable for allele-specific analyses. After filtering, 1,047 BAM files from 293 donors remained.

To ensure consistency in mapping and genome build alignment, BAM files were converted to FASTQ format and assessed with FastQC [43]. All FASTQ files had acceptable read quality (base quality >20) with no overrepresented sequences. Reads were remapped to the hg38 reference genome using STAR v2.7.10a [44] with parameters --outFilterType BySJout and --alignEndsType EndToEnd. Duplicate reads were removed using Picard’s MarkDuplicates. Heterozygous SNPs for each donor were extracted from matched WGS data, with VCF files converted from hg19 to hg38 using GATK’s LiftoverVCF tool [45]. Only heterozygous SNPs labeled “PASS” were retained for downstream analyses.

Allele-specific APA (asAPA) events were identified using the ASARP pipeline [11].Briefly, ASARP detects local allele-specific expression (ASE) by comparing allelic ratios between candidate SNPs (within alternative regions due to APA) and reference SNPs (outside APA regions). Global ASE was first filtered out to isolate local effects specific to 3’ UTR APA. Normalized expression values (NEVs) were calculated for annotated 3’ UTR regions using GENCODE annotations. These values were derived from the usage of 3’ UTR regions relative to constitutive exons, with the average read coverage of constitutive exons being normalized to a value of 1 and regions with coverage below that of the constitutive exons having a value between 0 and 1. Regions with NEV ≤ 0.8 were thus considered to exhibit APA. Testable genes for asAPA discovery were those that satisfy: (1) had APA patterns (NEV ≤ 0.8); (2) contained ≥1 heterozygous SNP in the extended region of the 3’ UTR; (3) the heterozygous SNP had ≥3 reads; and (4) had ≥1 control heterozygous SNP with ≥20 read coverage and an allelic ratio between 0.45 and 0.55 (indicating no large allelic imbalance) in a constitutively included region of the gene. The heterozygous SNP in the extension region was identified as an asAPA tag SNP if it showed significant allelic bias relative to the control SNP (Chi-square test, FDR < 0.05).

### Gene Ontology (GO) enrichment analysis

GO enrichment analysis was performed using Ensembl annotations and the biomaRt R package [46,47]. Each asAPA gene was matched to a background control gene selected from the set of genes testable by ASARP but not found to be significant. Control genes were required to match the asAPA gene in expression level and gene length (within 10%). To assess enrichment, we generated 10,000 random set of controls, each with the same number of genes as the set of asAPA genes, to create a null distribution. P-values for each GO term were calculated by comparing its frequency in the asAPA set against the null distribution. Statistical significance was determined using an empirical p value cutoff of 1 divided by the total number of GO terms tested. The same procedure was followed for analyzing the genes shortened in the FXS condition.

### Inference of putative functional asAPA SNPs with concordance analysis

To identify functionally relevant asAPA SNPs, we applied a concordance-based framework described by Amoah et al. (2021) [14]. This approach assumes that a truly functional SNP shows a consistent relationship between genotype and APA isoform usage across individuals. For each asAPA SNP, we calculated a concordance score for heterozygous candidate SNPs based on the SNP’s allelic ratio, or the proportion of reads carrying the reference allele (*R*_i_). Allelic imbalance was defined as *d*_i_ = |0.5 - *R*_i_|. Concordance scores were then derived from these values to quantify the alignment between observed allelic imbalance and candidate SNP genotype:

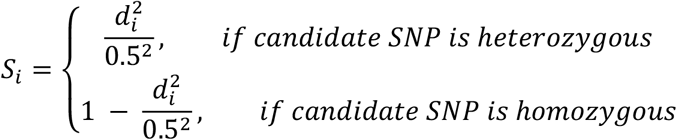

We applied this method to all common dbSNPs within the same gene as each asAPA SNP, generating a distribution of concordance scores across the population. APA-influencing SNPs are expected to produce distributions with either (i) a single peak near 1, or (ii) two peaks, one near 1 and another between 0 and 1, reflecting heterozygous individuals. A Gaussian mixture model was fitted to each distribution to identify these peaks. Significance was assessed by comparing observed peak heights to a background distribution generated from randomized concordance scores. SNPs were classified as putatively functional if they displayed significant peak enrichment and sufficient sample representation. Representation was considered adequate if ≥10% of tested individuals were heterozygous and the number of concordance values (*S*_i_) for the candidate-asAPA SNP pair exceeded 10% of the total sample size for a given brain region and condition.

### Enrichment of asAPA SNPs in RBP binding sites

To investigate the potential regulatory role of RBPs in asAPA, we analyzed ENCODE eCLIP-seq data from HepG2 and K562 cell lines [15], covering 127 RBPs. For each RBP, peak regions with at least four-fold enrichment over input controls were selected. Putatively functional asAPA SNPs were then tested for overlap with these enriched peaks.

To assess enrichment, we compared the number of asAPA genes harboring putatively functional SNPs in binding peaks of each RBP to a control set of 656 genes that were testable by ASARP but not found to be significant. We calculated fold enrichment as the ratio between the proportion of putatively functional asAPA genes bound by each RBP and the proportion of bound control genes. Bootstrapping was performed by sampling the same number of control genes as there are putatively functional asAPA genes 10,000 times to create a null distribution of binding proportions. A Gaussian distribution was fitted to the null, and a z-score and empirical p-value were computed for the observed enrichment. Multiple testing correction was applied using the Benjamini-Hochberg method [48]. RBPs with log2 fold change > log2(1.5) (∼0.58) and FDR < 0.05 were classified as significantly enriched beyond expected background levels.

### De novo motif identification using HOMER2

To identify sequence motifs potentially mediating allele-specific APA, we used HOMER2 [49], a *de novo* motif discovery tool designed to uncover overrepresented short motifs in input sequences relative to a background set. For each putatively functional asAPA SNP, we extracted an 11-nucleotide sequence centered on the SNP. We then conducted two complementary enrichment analyses via HOMER2: one using sequences harboring the higher expressed alleles as the input set and lower-expressed alleles as the background, and the other in reverse. Enriched motifs were then matched to known RNA-binding motifs from the RNA Bind-n-Seq (RBNS) dataset [16], requiring an exact 6-mer match. For *FMR1*, which recognizes shorter sequence elements, we additionally required an exact match to its known 4mer-binding motifs [17].

### Prediction of allele-specific RBP binding using DeepRiPe

To evaluate whether putatively functional asAPA SNPs alter RBP interactions, we applied DeepRiPe, a deep learning framework that predicts the impact of SNVs on RBP binding [50]. For each SNP, we calculated the absolute difference in predicted binding scores between the two alleles. As a negative control, we used an equal-sized set of SNPs testable by ASARP but not found to be significant. We then compared the distribution of binding score changes between asAPA SNPs and controls to assess whether asAPA SNPs were more likely to disrupt RBP interactions.

### Analysis of APA changes and FMRP binding in FXS

To test whether FMRP binding is associated with APA dysregulation in FXS, we analyzed RNA-seq data from postmortem frontal cortex samples of FXS and control individuals [22], using annotated polyadenylation sites from polyA_DB [23]. Reads were aligned to the hg38 genome with STAR, as described above. APA changes were quantified by averaging read coverage in the 400 nt window upstream of proximal and distal polyA sites (core and extension regions, respectively), separately for FXS and control groups. Sites with <400 bp separation were excluded, as subsequent analyses required regions to be 400 bp or greater. Directionality was determined from the log_2_ ratio of FXS-to-control coverage at proximal versus distal sites: transcripts were classified as shortened if the ratio exceeded 0.5 and lengthened if below −0.5.

To assess whether APA changes were associated with FMRP binding, we searched transcripts for known FMRP motifs (ACUG, ACUU, UGGA, AGGA) in the extension (proximal to distal APA site) and core regions (common 3’ UTR region). Regions <400 bp were excluded to ensure that each 1% sliding window across the region could contain a FMRP motif. Motif frequency was calculated using a sliding window spanning 1% of each region and counting the number of motifs within each. The frequency was then smoothed by averaging across ±5% of the flanking sequences. This enabled us to compare motif distributions near proximal and distal sites among transcripts classified as lengthened, shortened, or unchanged in FXS.

### Identification of asAPA SNPs in LD with GWAS SNPs

We assessed whether asAPA SNPs were linked to known disease-associated variants by comparing them to GWAS SNPs associated with ten neurodevelopmental and neurodegenerative disorders in the GWAS Catalog [1,24]. LD was evaluated using population reference panels from the EUR, AFR, EAS, and SAS groups. SNP pairs were considered in LD if they met the criteria R^2^ ≥ 0.8, D’ ≥ 0.9 and were located within 200kb of each other. To estimate the specificity of observed overlaps, we repeated the analysis using a matched background set of SNPs testable by ASARP but not found to be significant. To quantify enrichment, we calculated the Disease Relevance Ratio (DRR) for each disorder, defined as the proportion of asAPA SNPs in LD with GWAS loci divided by the proportion of matched control SNPs in LD with the same loci. A DRR >1 indicates enrichment of GWAS-linked variants among asAPA SNPs relative to background expectation.

### Definition of AD status

To determine AD status, we integrated multiple neuropathological criteria. Primary classification was based on the CERAD score, which stratifies brains as “no AD”, “possible AD”, “probable AD”, or “definite AD”. To refine these categories, we incorporated the Clinical Dementia Rating (CDR) and Braak neurofibrillary tangle stage, following prior recommendations [27] that assigned samples to four AD pathological levels: None, Low, Intermediate, or High. Samples were classified according to the following table:

| CDR | Braak |  |  | CERAD |
| --- | --- | --- | --- | --- |
|  | 0 - 2 | 3 - 4 | 5 - 6 |  |
| 0 | None | None | None | “No AD” |
| 1-2 | Low | Low | Low | “No AD” or “Possible AD” |
| 1-2 | Low | Intermediate | Intermediate | “Probable AD” or “Definite AD” |
| 3 | Low | Intermediate | Intermediate | Any |
| 4-5 | Low | Intermediate | Intermediate | “No AD” or “Possible AD” |
| 4-5 | Low | Intermediate | High | “Probable AD” or “Definite AD” |

For downstream analysis, we grouped None and Low pathology samples as controls, and Intermediate and High pathology samples as AD. This yielded 611 AD and 436 control samples for analysis.

### Differential asAPA detection using REDIT-LLR

To identify allele-specific APA events that differ between AD and control brains, we applied the REDIT-LLR statistical framework [28], adapted for allelic usage analysis. For each SNP *s* in sample *i* from group *g* (*g* ∈ {*AD, Control*}), we record the number of alternative-allele reads *Y*_*s,i,g*_ and the total read count n_*s,i,g*_. These counts were modeled as:

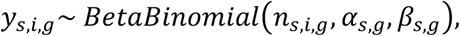

where *α*_*s,g*_ and *β*_*s,g*_ are group-specific shape parameters describing the distribution of true allelic usage for SNP *s* in group *g*.

For each SNP *s*, we compared a null model in which both groups share a single parameter set (*α*_*s,o*_, *β*_*s,o*_) to an alternative model where each group has its own parameters *α*_*s,AD*_, *β*_*s,AD*_ and *α*_*s,Ctrl*_, *β*_*s,Ctrl*_. Likelihood ratio statistics were calculated to evaluate whether allelic usage distributions differed between AD and control groups. A SNP was eligible for testing if it was identified as an asAPA SNP in either condition and had sufficient coverage (≥5 testable samples) in both AD and control groups. SNPs were classified as differentially used if they had a REDIT-LLR p-value below 0.05 and an absolute difference in average allelic usage between groups greater than 0.1. This was done to balance statistical significance with biological relevance.

To assess model adequacy, we evaluated each fitted (*α*_*s,g*_, *β*_*s,g*_) pair using a randomized probability integral transformation (PIT)-based Kolmogorov-Smirnov (KS) test [51,52]. For fitted parameters (*α*_*s,g*_, *β*_*s,g*_), we computed

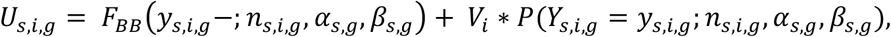

where *F*_*BB*_ is the beta-binomial cumulative distribution function and *F*_*BB*_ (*Y*_*s,i,g*_ −; *n*_*s,i,g*_, *α*_*s,i,g*_,*β*_*s,i,g*_) is the lower limit of *F*_*BB*_ at *Y*_*s,i,g*_, *V*_*i*_ is a random number from a uniform distribution on (0,1). Under a correctly specified model, *U*_*s,i,g*_ follows a *U*n*iform*(0,1) distribution. The KS test compared the empirical distribution of *U*_*s,i,g*_ to the expected uniform distribution, thereby evaluating how well the beta-binomial model fits the observed data. Exact KS p-values were obtained for each (*s, g*) pair and adjusted for multiple testing across all SNP-group pairs using the Benjamini-Hochberg procedure.

A total of 12 sites (among 2,558) in the control group and 8 sites (among 2,558) in the AD group had adjusted p < 0.05. Although these sites exhibited deviations from the beta-binomial fit, the majority were well-captured by the model, supporting its suitability for testing differences in allelic usage at polyadenylation-associated SNPs (Supplementary Figure 4A).

## Acknowledgements

We thank members of the Xiao laboratory for helpful discussions and comments on this work. This work was supported in part by grants from the National Institute of Health (R01AG075206, and R01AG078950).

## Competing interest statement

The authors declare no competing interests.

## Notes

### Competing Interest Statement

The authors have declared no competing interest.

