## Supplemental Figures for "Allele-specific alternative polyadenylation links noncoding genetic variation to Alzheimer’s disease risk"

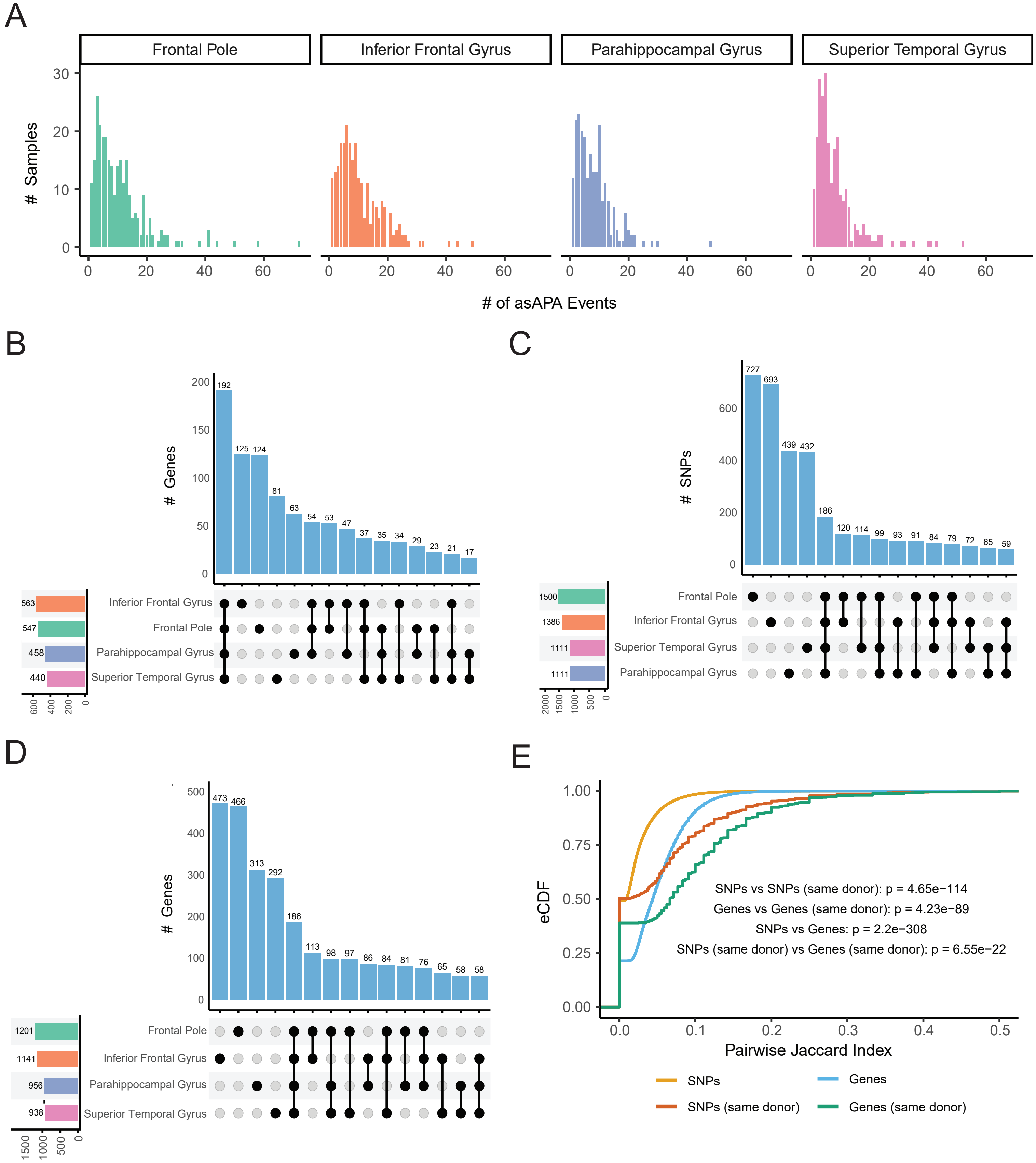


Supplementary Figure 1. Regional and inter-individual patterns of asAPA events in MSBB brain RNA-seq data. (A) Histogram showing the number of asAPA events per sample across brain regions. (B) Upset plot showing overlap of asAPA genes across the 4 brain regions where only those testable in all brain regions were included. (C) Upset plot showing the overlap of asAPA SNPs across the 4 brain regions. (D) Upset plot showing overlap of asAPA SNPs across the 4 brain regions where only those testable in all brain regions were included. (E) Empirical cumulative distribution function of pairwise Jaccard index similarity across any pair of brains or any pair of samples from the same donor brain for asAPA SNPs and genes. Jaccard index was calculated by taking the intersection over union across two brains (for SNPs, Genes) or by taking the intersection over the union across two samples (for same donor SNPs, same donor Genes). Kolmogorov-Smirnov test p values are shown comparing different groups.

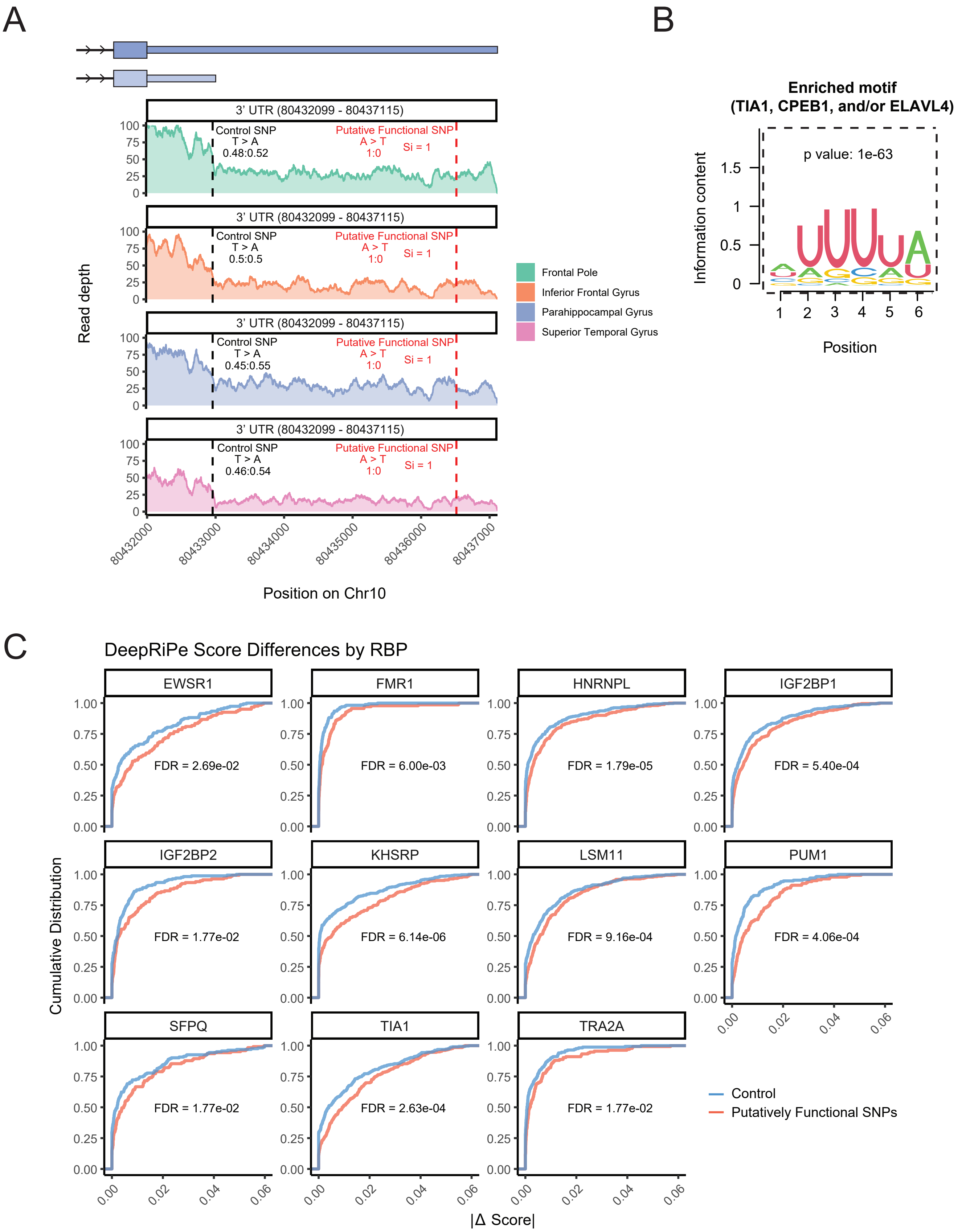


Supplementary Figure 2. Putatively functional asAPA SNPs and their impact on RBPs.
(A) Example of a putatively functional asAPA SNP in PRXL2A identified across brain region samples from distinct brain donors. The control SNP is labeled in the core 3' UTR region (black) and the putatively functional SNP is labeled in the extended 3'UTR region. The allelic ratio (reference:alternate) of both SNPs is displayed, and the concordance score (Si) is given for the putatively functional SNP (B) Representative motif identified by HOMER2 de novo motif discovery near putatively functional asAPA SNPs (higher expressed alleles). The motif matches known binding preferences of RBPs including TIA1, CPEB1, and ELAVL4, based on RBNS motif database comparisons. Hypergeometric test p-value comparing motif frequency in higher expressed alleles to lower expressed alleles is shown (C) Violin plots showing the absolute difference in DeepRiPe-predicted RBP binding scores between alleles of putatively functional asAPA SNPs versus control SNPs, for 11 RBPs. Putatively functional SNPs (red) consistently exhibit greater predicted allele-specific effects on RBP binding. Wilcoxon rank-sum tests * p<0.05; ** p<0.01; ***p<0.001; ****p<0.0001.

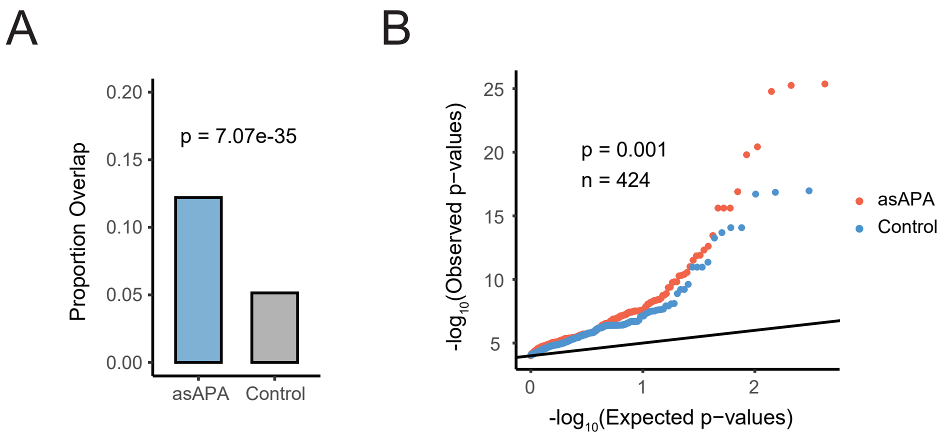


Supplementary Figure 3. Overlap of asAPA SNPs with aQTL signals.
(A) Bar plot comparing the proportion of SNPs overlapping known aQTLs. All significant asAPA SNPs vs. control SNPs, where the control SNP set was defined as the set of SNPs tested by the ASARP pipeline but not found to be significant. p-values were calculated using a Chi-square test. (B) QQ plot comparing aQTL p-value distributions for significant asAPA SNPs (red) vs. matched control SNPs (blue). p-value derived from a Kolmogorov-Smirnov test.

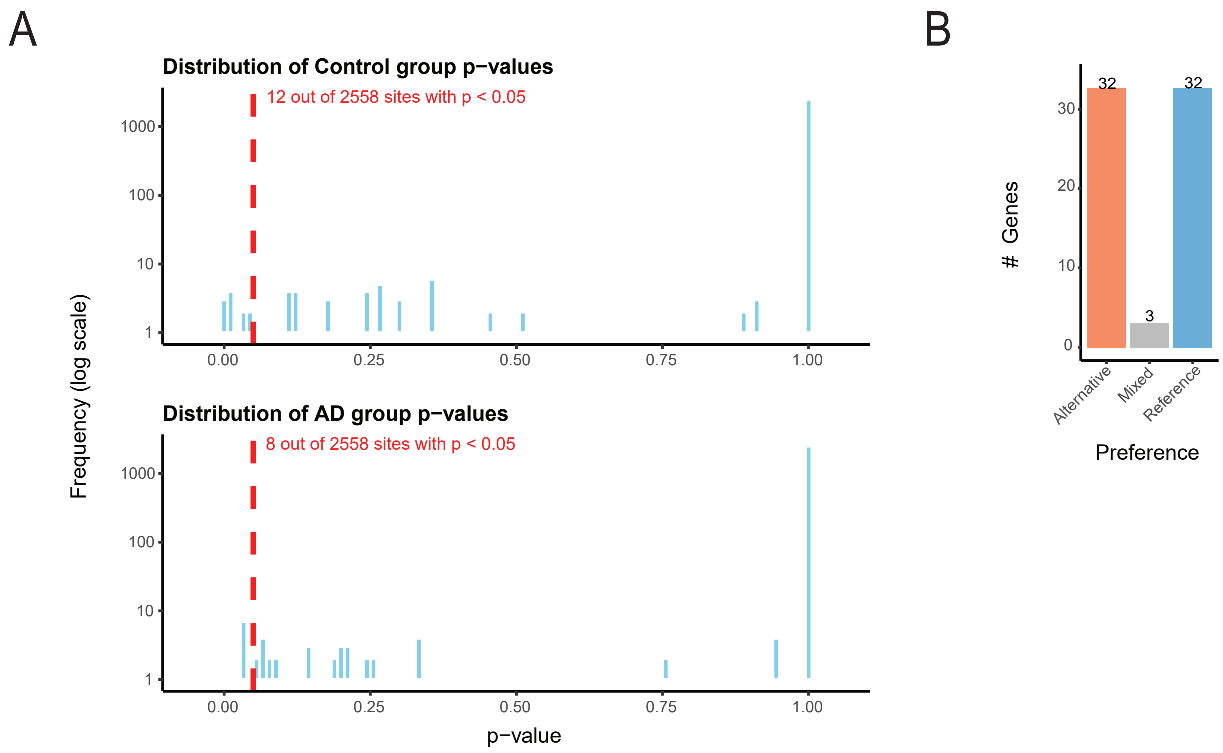


Supplementary Figure 4. Model validation and allelic preference patterns in AD
(A) Distribution of p-values from randomized posterior predictive checks using the PIT-based Kolmogorov-Smirnov (KS) test in the control (top) and AD (bottom) groups (Methods). The majority of sites do not show significant deviation from the beta-binomial model, supporting the suitability of the REDIT-LLR framework for modeling allelic APA. (B) Allelic preference in genes with significant differential asAPA in AD. Preference categories are defined based on whether the reference or alternative allele showed higher usage in AD. “Mixed” indicates genes with inconsistent allele preference across brain regions.
